# *Legionella pneumophila* CRISPR-Cas suggests recurrent encounters with *Gokushovirinae*

**DOI:** 10.1101/2021.03.08.434514

**Authors:** Shayna R. Deecker, Malene L. Urbanus, Beth Nicholson, Alexander W. Ensminger

**Affiliations:** Department of Biochemistry, University of Toronto, Toronto, Ontario, Canada; Department of Molecular Genetics, University of Toronto, Toronto, Ontario, Canada

## Abstract

*Legionella pneumophila* is a ubiquitous freshwater pathogen and the causative agent of Legionnaires’ disease. This pathogen and its ability to cause disease is closely tied to its environmental encounters. From phagocytic protists, *L. pneumophila* has “learned” how to avoid predation and exploit conserved eukaryotic processes to establish an intracellular replicative niche. Legionnaires’ disease is a product of these evolutionary pressures as *L. pneumophila* uses the same molecular mechanisms to replicate in grazing protists and in macrophages of the human lung. *L. pneumophila* growth within protists also provides a refuge from desiccation, disinfection, and other remediation strategies. One outstanding question has been whether this protection extends to phages. *L. pneumophila* isolates are remarkably devoid of prophages and to date no *Legionella* phages have been identified. Nevertheless, many *L. pneumophila* isolates maintain active CRISPR-Cas defenses. So far, the only known target of these systems has been an episomal element that we previously named *Legionella* Mobile Element-1 (LME-1). In this study, we have identified over 150 CRISPR-Cas systems across 600 isolates, to establish the clearest picture yet of *L. pneumophila*’s adaptive defenses. By leveraging the sequence of 1,500 unique spacers, we can make two main conclusions: current data argue against CRISPR-Cas targeted integrative elements beyond LME-1 and the heretofore “missing” *L. pneumophila* phages are most likely lytic gokushoviruses.

**IMPORTANCE:** The causative agent of Legionnaires’ disease, an often-fatal pneumonia, is an intracellular bacterium, *Legionella pneumophila*, that normally grows inside amoebae and other freshwater protists. Unfortunately for us, this has two major consequences: the bacterium can take what it has learned in amoebae and use similar strategies to grow inside our lungs; and these amoebae can protect *Legionella* from various forms of chemical and physical disinfection regimes. *Legionella* are ubiquitous in the environment and frequently found in man-made water systems. Understanding the challenges to *Legionella* survival before it reaches the human lung is critical to preventing disease.

We have leveraged our earlier discovery that *L. pneumophila* CRISPR-Cas systems are active and adaptive – meaning that they respond to contemporary threats encountered in the environment. In this way, CRISPR arrays can be considered genomic diaries of past encounters, with spacer sequences used to identify elements that may impinge on the pathogen’s survival. One outstanding question in the field is whether *L. pneumophila* is susceptible to phage, given the presumptive protection provided by intracellular replication within its eukaryotic hosts. In this work, we use CRISPR spacer sequences to suggest that the heretofore “missing” *L. pneumophila* phage are most likely lytic gokushoviruses. Such information is critical to the long-term goal of developing of new strategies for preventing colonization of our water systems by *Legionella* and subsequent human exposure to the pathogen.

## INTRODUCTION

*Legionella pneumophila* is a Gram-negative, intracellular bacterium that is ubiquitous in freshwater environments (1-3), where it replicates within a wide range of protist hosts (4). If a contaminated water source becomes aerosolized and inhaled, *L. pneumophila* can infect human lung macrophages and cause a severe pneumonia known as Legionnaires’ disease (5, 6). Replication in the accidental human host uses similar molecular strategies to those used to infect protists (7). As humans are an evolutionary dead end to the pathogen (8), understanding how *L. pneumophila* is able to persist and replicate in environmental reservoirs is critical to limiting its ability to cause human disease.

Protozoan hosts not only serve as a replicative niche to *L. pneumophila* (1, 4), but also provide protection from desiccation (9), temperature changes (10, 11) and disinfectants (10-13). One outstanding question is whether these hosts also protect *L. pneumophila* from predation by foreign genetic elements, such as viral bacteriophages (phages). Notably, phages for the obligate intracellular pathogens *Chlamydia psittaci* and *Chlamydia pecorum* have been described (14-17), raising the possibility that phages may access *L. pneumophila* even within the protection of the host.

The first report of *Legionella* infection by lytic phages isolated from freshwater samples is equal parts promise and obfuscation (18). Preliminary analysis suggested these phages were members of the *Myoviridae* family (18), but inefficient phage enrichment prevented the preservation of stocks for validation and further study. Around the same time, another group reported visualization of temperate *Legionella* phage, but this occurred in a well-characterized, fully sequenced type strain with no potential to generate phage particles (19). Now a decade later, no subsequent studies have confirmed either laboratory’s findings or identified any other type of *L. pneumophila* phage. Lysogeny also seems uncommon in the *Legionella* genus as the only known “prophage-like” elements described to date are *Legionella* mobile element-1 (LME-1), which has been proposed to descend from an ancestral phage (20, 21), and a putative prophage in one *Legionella micdadei* isolate (22).

Many *L. pneumophila* isolates maintain active and adaptive CRISPR-Cas systems (21, 23, 24), providing another avenue by which to explore the species’ relationship with phage. Because *L. pneumophila* CRISPR-Cas protects against foreign threats, the CRISPR array in each *L. pneumophila* genome serves as a sequence-based diary of past environmental encounters. When a bacterium encounters a foreign element (e.g. a plasmid or a phage), a short DNA sequence (spacer) can be obtained and incorporated into a CRISPR array (25-28). The transcribed CRISPR RNA forms a complex with associated *cas* genes, and uses complementary base-pairing and endonuclease activity to cleave the foreign element and protect the bacterium (29-32). Given the sequence-based functionality of CRISPR-Cas systems, sequence similarity can be applied to identify targets of these systems for individual bacterial species (21, 33) and in a global context (34-37).

We previously used spacer sequence similarity to identify the first known target of *L. pneumophila* CRISPR-Cas systems, a mobile genetic element known as LME-1 (21). In the same study, we showed that LME-1 restricts host range and that *L. pneumophila* CRISPR-Cas successfully defends against transfer of the element between strains (21). However, we found that only ∼3% of *L. pneumophila* spacers match to LME-1 (21), leaving the vast majority of spacers without defined targets. In this study we have expanded our analyses to include a collection of over 600 *L. pneumophila* isolates and 1,500 unique spacers. Leveraging this expanded dataset, we identify only two recurrent targets of *L. pneumophila* CRISPR-Cas: LME-1 and the *Gokushovirinae* family of phages.

## MATERIALS AND METHODS

### *Legionella* genomes used in this study

*Legionella pneumophila* genomes (draft and completed) were downloaded from the European Nucleotide Archive (at: https://www.ebi.ac.uk/ena/browser/home) (38) and from NCBI (at: https://www.ncbi.nlm.nih.gov/) (39). A complete list of accession numbers can be found in Table S1.

Some target sequence data were produced by the US Department of Energy Joint Genome Institute (at: http://www.jgi.doe.gov/) in collaboration with the user community (Table S7).

### Prophage identification

The downloaded *L. pneumophila* genomes were annotated with Prokka (v.1.14.6) (40) and the resulting GenBank formatted files were analyzed using PhiSpy (41) with the default settings. The original nucleotide FASTA files were analyzed using VirSorter (42) with the default settings.

CsrT amino acid sequences from mobile elements (43) identified by PhiSpy and VirSorter were extracted and aligned with MUSCLE (44) and a tree was generated using FastTree (v 2.1.11) with the default settings (45). The tree was rooted to group I, Trb (43), and visualized using the Interactive Tree of Life (iTOL) web server (46). Reference sequences were annotated as per Abbott and colleagues (43).

The 29 nucleotide LME-1 attachment site (GTCTGATTATCAAAATAATCAGACTTAAT) (21) was queried by BLASTn against *L. pneumophila* genomes using the following parameters: -gapopen 10 -gapextend 2 -reward 1 -penalty -1 -evalue 1 -word_size 7 (47).

### Core genome identification across LME-1 variants

The FASTA sequences for unique LME-1 variants were subjected to an OrthoMCL analysis to look at the overlap in gene content. Briefly, the sequences were annotated using Prodigal (48), then were analyzed using OrthoMCL (v.2.0.9) with the default settings (49). The resulting clusters were queried by BLASTp using the NCBI viral database (taxid:10239) (47) and using HHpred (at: https://toolkit.tuebingen.mpg.de/tools/hhpred) (50) against “UniProt-SwissProt-viral70_23_Aug_2020” using default settings to determine if any had sequence similarity to known phage genes.

Core genomes for each LME-1 variant, containing only genes present within all the variants, were aligned using MUSCLE (44) and a tree was generated using FastTree (v 2.1.11) using the default settings and the GTR model (45). The tree was visualized with ggtree (v. 2.2.4) (51), and the orthologous group presence/absence map was plotted with ggplot2 (v. 3.3.3) (52).

### CRISPR-Cas identification and spacer cataloguing

CRISPRCasFinder (53) was used to detect putative CRISPR-Cas systems in *L. pneumophila* genomes (Table S1) and CRISPRDetect (54) was used to determine the transcriptional direction of each CRISPR array. Spacers were extracted from each array and compiled to form spacer libraries. Redundant spacers were removed from the libraries to create a unique spacer library for downstream analyses. Rarefaction curves for the spacer library of each CRISPR-Cas sub-type were generated using Analytic Rarefaction (v. 2.2.1) using the default parameters (available from https://strata.uga.edu/software/index.html). The curves were plotted using ggplot2 (v. 3.3.2) (52).

### CRISPR-Cas target search

A pipeline to search for targets of *L. pneumophila* CRISPR-Cas was written as a Unix shell script. *L. pneumophila* genomes were first searched for putative type I-C, I-F and II-B CRISPR arrays using the repeat structure as a guide. If a CRISPR array was detected, it was subsequently masked to minimize false positive hits to the input genomes. A BLAST database was then constructed containing the *Legionella* genomes with masked CRISPR arrays to look for integrated genetic elements (see Table 1). The unique spacer library (see above) was queried against the BLAST database (BLASTn parameters: -gapopen 10 -gapextend 2 -reward 1 –penalty -1 -evalue 0.01 -word_size 7) (47). The BLAST search was done twice, once for each strand. The BLAST output was then processed to score each hit. The scoring metric subtracted the length of the alignment between a spacer and a hit from the length of the spacer, then added the number of mismatches between the two sequences. For example, an alignment length of 30, a spacer length of 32 and 0 mismatches would generate a score of 2. This was done to take both alignment length and number of mismatches into account when ordering the hits for downstream analyses (for example, so a hit with 0 mismatches and a short alignment length would not be ranked higher than a hit with 2 mismatches but an alignment length across the entire spacer).

**Table 1.** Target databases. The databases used for spacer target searches for integrated elements and exogenous elements.

| Database Name | Target Type | Date Downloaded | Number of Genomes, Reads or Contigs | Genome Completion | Reference |
| --- | --- | --- | --- | --- | --- |
| European Nucleotide Archive - *Legionella pneumophila* | Integrated elements | 2019-01-09 | 519 | Draft | 38 |
| NCBI - *Legionella pneumophila* | Integrated elements | 2019-04-08 | 83 | Complete | 39 |
| Sequence Read Archive - *Legionella pneumophila* | Integrated elements | 2019-04-08 | 9 | Raw reads | 21 |
| DataONE Dash Plasmids | Exogenous elements | 2019-04-05 | 10,892 | Complete | 63 |
| NCBI - *Microviridae* | Exogenous elements | 2019-12-02 | 2,864 | Complete & draft | 39 |
| NCBI - Viruses | Exogenous elements | 2019-04-08 | 28,652 | Complete | 39 |
| IMG/VR | Exogenous elements | 2020-01-29 | 734,969 | Metagenomic | 66, 67 |
| *Microviridae* associated contigs from IMG/M | Exogenous elements | 2020-02-04 | 14,724 | Metagenomic | 68 |
| *Microviridae* associated contigs from marine sources | Exogenous elements | 2020-07-30 | 12,775 | Metagenomic | 64, 65 |
| Oxford Nanopore read from an aquifer | Exogenous elements | 2020-06-25 | 2,380,279 | Metagenomic | 69 |

After scoring, the BLAST output was converted into a BED format. During this step, the putative target sequences were extended to account for alignment length, followed by the addition of a 3 nucleotide extension on either end of the sequence for downstream protospacer adjacent motif (PAM) filtering. SAMtools (55) and BEDTools (56) were used to extract the putative target sequences and their flanking regions from the input genomes. Finally, the pipeline searched for canonical PAMs for the type I-C, I-F and II-B systems in the flanking region of the putative target sequence. Although our pipeline searched for type I-C, I-F and II-B PAMs, we considered this secondary information and were intentionally PAM agnostic when assigning spacers to target groups to avoid missing potential targets that have since escaped through PAM mutations or those targeted by non-canonical PAMs. Analysis of the PAM sequences present in the target sequences showed that many target sequences did contain the canonical PAM (Figure S3).

The pipeline was also used to search for exogenous threats to the bacterium in viral, plasmid and metagenomic sequences and run as described above (data sets listed in Table 1). The output from the pipeline can be found in Table S6 for complete/draft genomes, and Table S7 for metagenomic data. CRISPR arrays containing spacers with ascribed targets were visualized with CRISPRStudio (57). PAM sequence logos for the hits were plotted using ggseqlogo (v 0.1) (58).

### Major capsid protein phylogeny of targeted *Microviridae* genomes

A phylogeny of *Microviridae* major capsid proteins was generated using the amino acid sequences from 3014 genomes (Table S8). The amino acid sequences were aligned using MUSCLE (44) and the tree was generated with FastTree (v 2.1.11) using the default settings (45). The resulting tree was rooted to the *Bullavirinae* clade and visualized using the iTOL web server (46). The sub-family designations for each phage based on the literature were plotted as a colour-coded ring circling the tree (Table S8).

### Sequence conservation and spacer mapping for repeat *L. pneumophila* CRISPR-Cas targets

Sequence conservation analysis and spacer mapping were performed on the *Microviridae* genomes that contained a top-hit target for a spacer that met the following stringency criteria: 5 or fewer mismatches with its protospacer, a canonical PAM for its CRISPR subtype, and no mismatches in the seed sequence region (positions 1-5, 7, 8). The VP1 (major capsid protein), VP2 (DNA pilot protein) and VP4 (replication initiation protein) genes were extracted from these genomes using Geneious Prime 2021 (available from:https://www.geneious.com/prime/), and the sequences for each gene were aligned using the MUSCLE plug-in (44). The alignment, consensus sequence, sequence identity profiles and spacer location for each gene were plotted using GViz (v. 1.32.0) (59), binning the sequence conservation with a sliding window of 5.

## Code availability

The script used to search for targets of *L. pneumophila* CRISPR-Cas systems is available on GitHub at https://github.com/EnsmingerLab/LegionellaCRISPRTargetSearch.

## RESULTS

### *Legionella* integrated elements identified by prophage prediction

The only target of *L. pneumophila* CRISPR-Cas identified to date is an integrated mobile element, LME-1 (21). It has been previously reported that LME-1 contains a number of predicted proteins of phage origin (20, 21), so we decided to revisit whether variants of LME-1 or other integrated *L. pneumophila* phage-like elements could be identified using established prophage prediction tools. We subjected a set of over 600 publicly available *L. pneumophila* genomes to analysis by two complementary prophage prediction programs, PhiSpy (41) and VirSorter (42) (Table S1), which use different methods to identify potential phage-like sequences. PhiSpy predictions are based on a variety of metrics such as AT/GC skew, phage sequence signatures, strand directionality and the presence of potential attachment sites (41). VirSorter identifies viral sequences by similarity to phage proteins (Hmmsearch and BlastP) and metrics associated with virus-like genome structures, such as the presence of hallmark viral genes, enrichment in uncharacterized genes and depletion in strand switching between consecutive genes (42).

Of 35 predictions that overlapped between VirSorter and PhiSpy, 7 were LME-1 variants -flanked by the previously described LME-1 attachment site and containing nearly identical gene content (Table S2). Each of the remaining 28 predicted regions contained *csrT*, a regulatory gene which is co-inherited with type IV secretion systems from *Legionella* integrative conjugative elements (ICEs) (43) (Table S2). By CsrT phylogeny, these ICEs all fall into the previously described group IV Lgi elements (Figure S1) (43).

To determine if any additional LME-1 variants might have been missed by our analyses, we searched each *L. pneumophila* genome for a signature of LME-1 integration: the presence of two LME-1 attachment (*att*) sites. This analysis identified 8 *att*-flanked insertions of >29 kb (Table S3). These integrants corresponded to all the previously identified LME-1 variants above, along with one additional LME-1 variant missed by PhiSpy. We also identified 24 instances of a short, 469 bp insertion at the LME-1 *att* site that shares no sequence similarity to LME-1, phages, or any other known sequence (Table S3). Unlike LME-1 insertions, the second *att* site for these instances contain several substitutions and insertions, suggesting against a contemporary integration event.

We next examined this expanded set of LME-1 variants for differences in gene content and sequence similarity to phage genes. Based on core genome phylogeny, the LME-1s segregate into 5 distinct variants with a set of 32 core genes and 17 accessory genes (Figure 1, Table S4). We examined each LME-1 predicted gene product for similarity to phage proteins using HHpred and BlastP. Strikingly, 27/32 of the core LME-1 genes share similarity to phage proteins (and 6/17 accessory genes). The preponderance of phage-like genes in the LME-1 core (including key phage structural proteins) clearly points to a phage origin (20, 21). It also raises the intriguing possibility that these sequences may generate phage-like particles under one or more heretofore untested experimental conditions.

**Figure 1.**
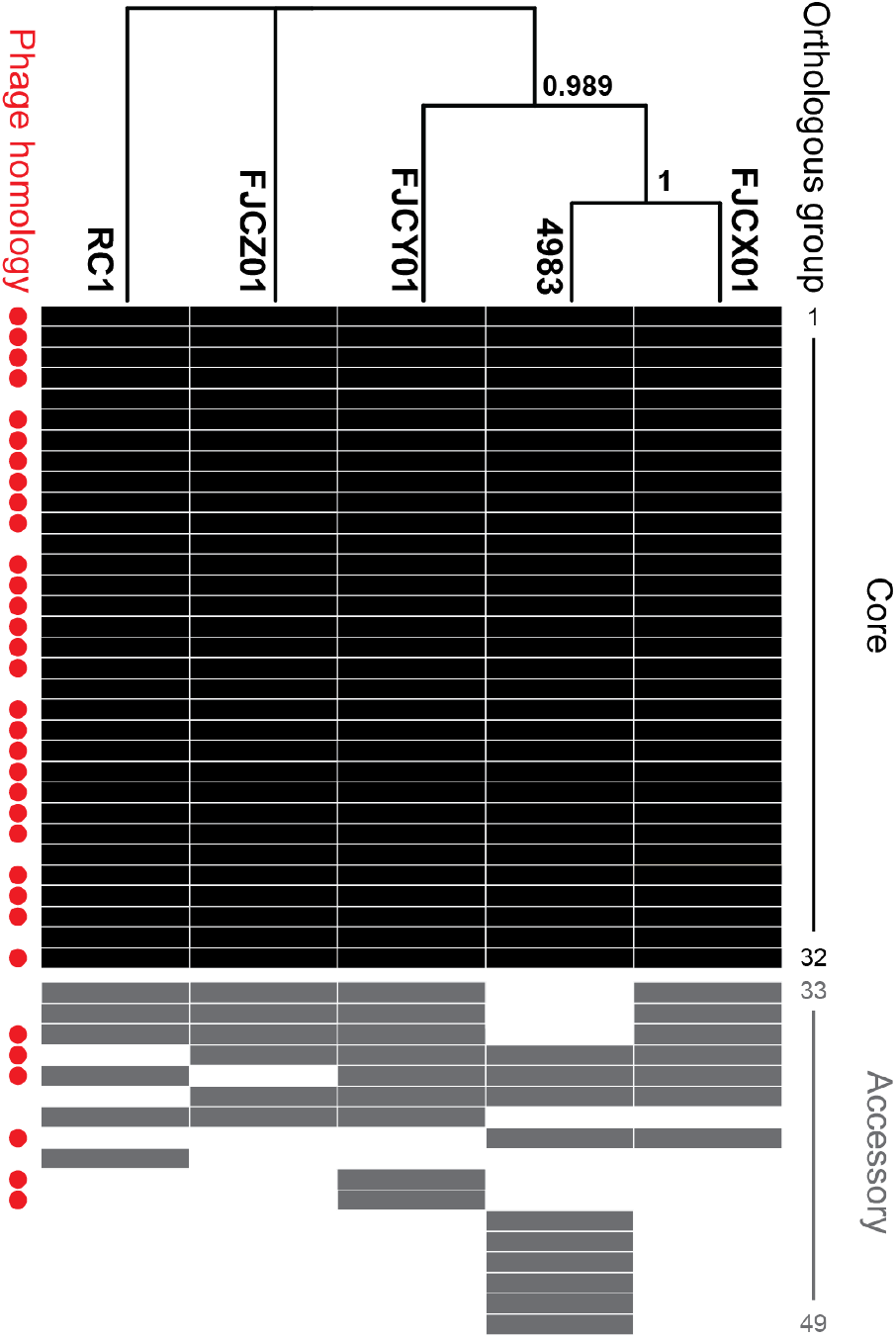
Core orthologous genes in LME-1 enriched for sequence similarity to known phage genes. Orthologous groups for LME-1 genes were determined using OrthoMCL. The core LME-1 genomes were aligned with MUSCLE; a phylogeny was constructed using FastTree and subsequently visualized in ggtree. Orthologous group presence/absence in each LME-1 genome was visualized with ggplot2. Red dots adjacent to each orthologous group indicate sequence similarity to known phage genes via BLASTp and/or HHPred predictions (Table S4).

### Cataloging the scope of *L. pneumophila* CRISPR-Cas defenses

Based on our analysis of 18 *L. pneumophila* isolates, we previously identified LME-1 as the only recurrent target of *L. pneumophila* CRISPR-Cas (21). Leveraging the availability of a drastically expanded genome set, we revisited our analysis of *L. pneumophila* CRISPR-Cas to determine if one or more additional targets might emerge. To begin, we systematically searched for CRISPR-Cas systems in over 600 *L. pneumophila* isolates (Table S1, Figure 2A) using CRISPRCasFinder (53) and CRISPRDetect (54). Combined with previously identified systems (21, 24, 60, 61), we identified a total of 13 isolates with type I-C CRISPR systems, 47 with type I-F systems and 108 with type II-B systems. These systems contain a total of 1589 unique spacers (Table S5), a four-fold increase relative to our earlier search for targets of *L. pneumophila* CRISPR-Cas (21). Nevertheless, rarefaction curves clearly show the continued value of further sequencing, especially for genomes containing type I-C systems (Figure 2B). This is consistent with our previous observations that at least two of these systems (I-C and I-F) are adaptive (21), suggesting that additional sampling of spacer sequences will continue to reveal new encounters between *L. pneumophila* and its invaders.

**Figure 2.**
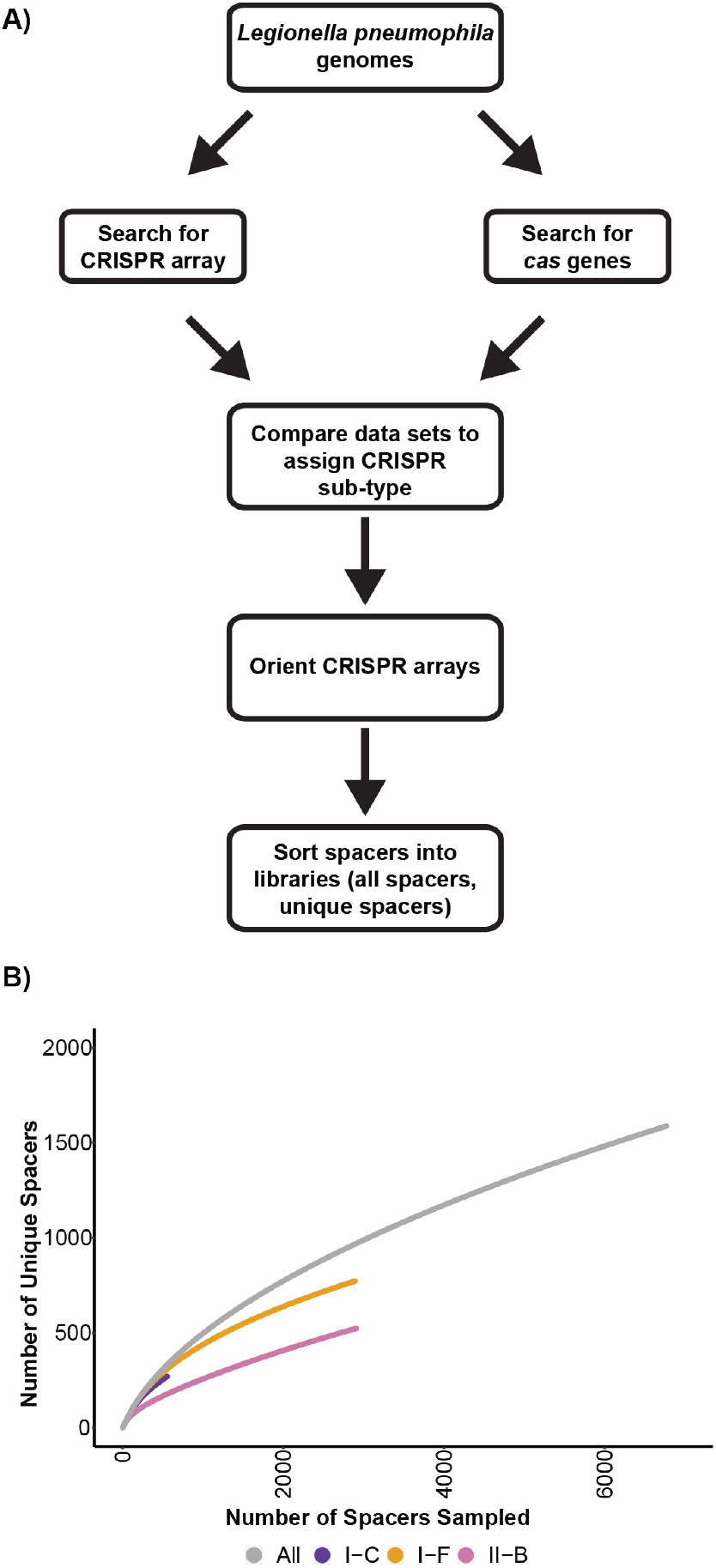
A catalogue of *L. pneumophila* CRISPR-Cas systems and their spacers. **A)** A schematic of the CRISPR-Cas system discovery pipeline used in this work. **B)** Rarefaction curves for each CRISPR-Cas sub-type in *L. pneumophila* suggest none of these sub-types were at saturation for unique spacer counts.

### LME-1 is the only integrative element targeted by *Legionella* CRISPR-Cas systems

With our expanded spacer library, we next sought to identify the matching protospacer sequences targeted by these systems. Our previous analysis, limited in both spacer sequences (queries) and *L. pneumophila* genomes (to search), nevertheless identified two variants of a LME-1 as frequent targets of all 3 types of *L. pneumophila* CRISPR-Cas (21). Backed by our prophage analysis above, we sought to determine if LME-1 remained uniquely targeted relative to other integrated genomic elements. We queried our expanded spacer set against a local BLAST database of over 600 *L. pneumophila* genomic sequences to search for any integrated elements that overlap with our prophage analysis or may have been missed (Table 1, Figure 3A) (38, 39).

**Figure 3.**
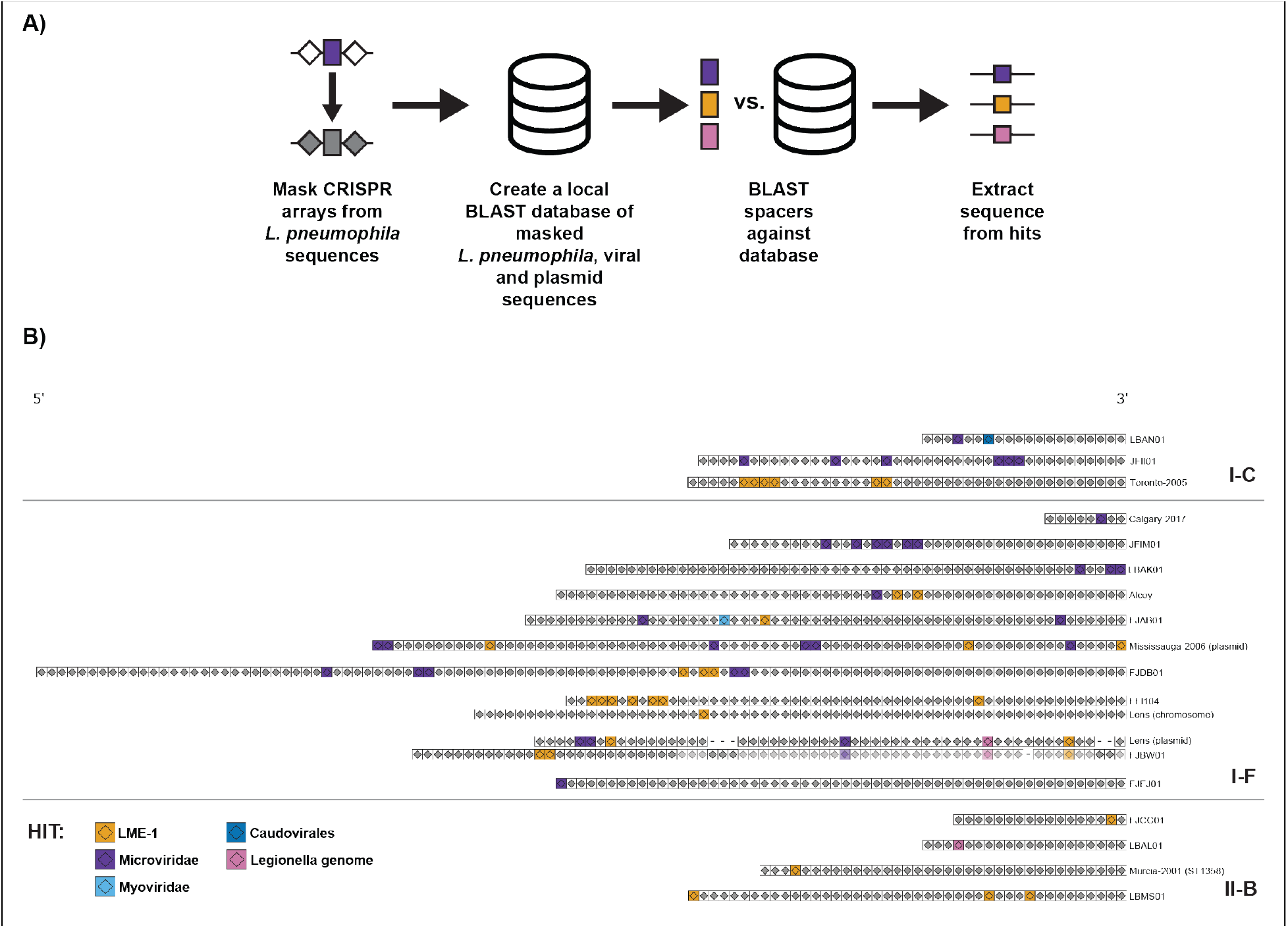
*L. pneumophila* CRISPR-Cas systems repeatedly target LME-1 and *Microviridae* phages. **A)** A Unix shell script was written to search for targets of *L. pneumophila* CRISPR-Cas. This pipeline masked putative type I-C, I-F and II-B CRISPR arrays in input genomes before creating a local BLAST database. It then queried the unique spacer library against this database using BLAST and extracted putative target sequences for downstream analyses. **B)** *L. pneumophila* CRISPR arrays possessing spacers with ascribed targets were visualized with CRISPRStudio; greyed boxes represent spacers without ascribed targets, while coloured boxes represent spacers with ascribed targets. Related arrays are grouped together, with spacer loss shown as dashes and shared spacers between those arrays shown in a lighter opacity.

Even within the expanded spacer catalog, LME-1 remains the only integrated target of *L. pneumophila* CRISPR-Cas. Thirty-two spacers target the element, representing approximately 2% of the unique spacer library (Table S6). LME-1 targeting spacers are present in 12 distinct CRISPR-Cas arrays, covering all three CRISPR-Cas sub-types (Table 2, Figure 3B). Within individual arrays, we observe multiple instances of adjacent LME-1 targeting spacers, consistent with repeat exposure and primed spacer acquisition (28) (Figure 3B). Of the 32 spacers that target LME-1, the vast majority (28) target all variants.

**Table 2.** CRISPR spacer distribution. A breakdown of the number of unique spacers for each CRISPR-Cas subtype, the number of spacers with identified targets and by target class.

| CRISPR-Cas Subtype | Type I-C | Type I-F | Type II-B |
| --- | --- | --- | --- |
| Number of Unique Spacers | 270 | 774 | 545 |
| Number of Spacers with Target | 13 | 52 | 6 |
| Percent of Library with Target | 4.81 | 6.72 | 1.10 |
| Number of Spacers Targeting *L. pneumophila* genome: LME-1 | 6 | 21 | 5 |
| Number of Spacers Targeting *Microviridae* | 6 | 29 | 0 |
| Number of Spacers Targeting *L. pneumophila* genome: other | 0 | 1 | 1 |
| Number of Spacers Targeting *Myoviridae* | 0 | 1 | 0 |
| Number of Spacers Targeting *Caudovirales* | 1 | 0 | 0 |

While two additional spacers mapped back to other *L. pneumophila* genomic targets (Table 2, Figure 3B, Table S6), neither fall within regions predicted by PhiSpy or VirSorter. Close inspection of the genomic regions surrounding these singleton hits does not suggest the presence of a mobile element or prophage based on the presence of phage-like proteins or %GC content. Notably absent from the collection of spacer hits are sequences matching the 5 endogenous plasmids (Paris (61), Lorraine (61), Lens (60, 61), Mississauga-2006 (21) and OLDA (62)) previously described in *L. pneumophila*. Likewise, we observe no matches between spacers and 22 additional endogenous *L. pneumophila* plasmids deposited into NCBI (Table S1) or even a larger, general database of plasmid sequences (63) (Table 1). Somewhat surprisingly, LME-1 remains the only integrative element (prophage-like or otherwise) targeted by *L. pneumophila* CRISPR-Cas.

### *L. pneumophila* CRISPR-Cas targets phages

The observation that LME-1 is the only prophage-like element targeted by CRISPR-Cas argues against widespread lysogeny in *L. pneumophila* and is consistent with prior observations that the *Legionella* genus is remarkably prophage-poor (22). The question remains, however, as to whether one or more lytic phages might pose an exogenous threat to *L. pneumophila* survival. To look for evidence of past *L. pneumophila*-phage encounters, we used the unique spacer library to query an extensive database of viral and phage sequences with over 28,000 viral genomes from NCBI (Table 1) (39). Within this collection, only one class of targets presented a signature similar to that of LME-1 (both in the number of unique spacers and the diversity of systems involved). Phages from the *Microviridae* family were, like LME-1, frequent targets of *L. pneumophila* CRISPR-Cas: targeted by 28 spacers across 11 distinct arrays (Table 2, Figure 3B, Table S6). Many of these spacers are found near the 5’ end of the array, suggesting a recent acquisition event as this is where new spacers are generally acquired (28) (Figure 3B). As with LME-1, several instances of adjacent *Microviridae*-targeting spacers can be found within individual arrays – again suggestive of multiple exposures and consistent with primed spacer acquisition (28) (Figure 3B).

Since many of the targeted *Microviridae* genomes were assembled from metagenomic reads and subsequently deposited into NCBI, we reasoned that there could be additional targets of *L. pneumophila* CRISPR-Cas present in metagenomic data. As such, we next searched for spacer hits across several metagenomic datasets (Table 1) (64-69). These analyses identified 7 additional spacers that targeted contigs of likely *Microviridae* origin, along with one additional spacer that targeted a contig of likely *Caudovirales* origin (Table 2, Figure 3B, Table S7).

In total, 35 *L. pneumophila* spacers target sequences of *Microviridae* origin: 25 spacers target the conserved major capsid protein VP1, 4 target the pilot protein VP2, and 5 target the DNA replication protein VP4. One spacer targets a non-coding region (Table 3). As with the spacers against LME-1, *Microviridae*-targeting spacers come from a genetically and geographically diverse set of *L. pneumophila* isolates. As with LME-1, this suggests that one or more *Microviridae* phage may represent a commonly encountered threat to *L. pneumophila* in the environment. The apparent frequency of these exposures, along with the lack of repeat spacers against any additional class of phage, significantly narrows the hunt for *L. pneumophila* phages.

**Table 3.**
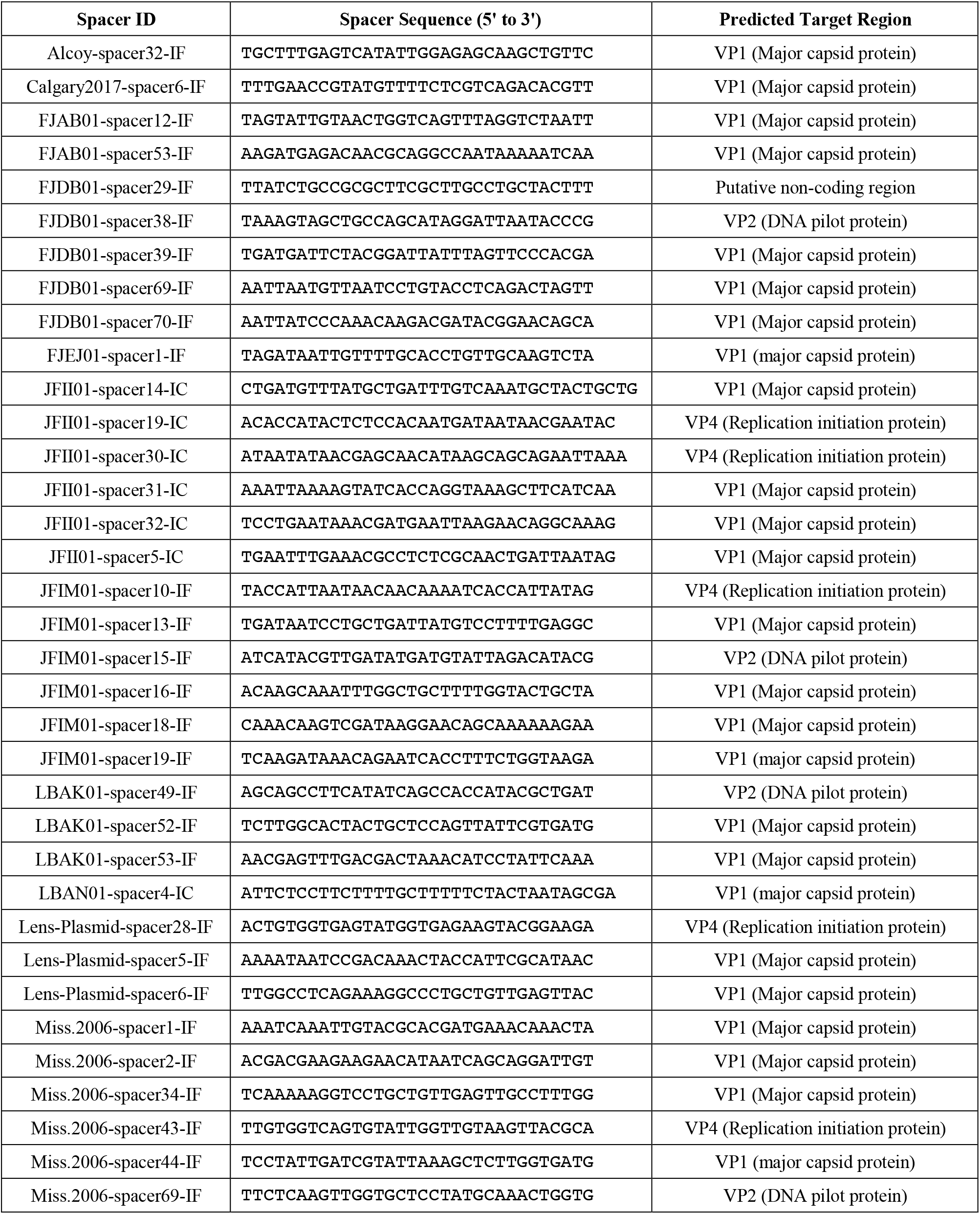
List of *Microviridae* target regions. The name, sequence and predicted target region for the unique spacers targeting *Microviridae*.

### L. pneumophila phages most likely belong to *Gokushovirinae*

The *Microviridae* family consists of the recognized subfamilies *Bullavirinae*, which includes well-characterized members such as phi-X174, alpha3 and G4 (70), and *Gokushovirinae*, first characterized as phages of *Chlamydia* (14-17), *Spiroplasma* (71-74) and *Bdellovibrio* (75). Several additional subfamilies have been proposed in recent years including: *Pichovirinae* (76), *Alpavirinae* (77), *Sukshmavirinae* (78), Dragonfly-associated Microvirus “Group D” (78, 79), *Parabacteroidetes* group based on *Parabacteroidetes* prophages (80, 81) and Pequeñovirus (80, 82).

To determine if we could further narrow the targets of *L. pneumophila* CRISPR-Cas to one or more of these subfamilies, we first generated a major capsid (VP1) phylogeny with all 3000 available *Microviridae* genomes (Figure 4, Table S8). We also included 14 spacer-targeted metagenomic contigs (identified above) with overlaps of 99-135 bp on the ends, as this overlap suggests these to be complete, circularizable *Microviridae* genomes (Table S7). We then indicated the top target for each spacer onto its corresponding phage genome on the tree. Strikingly, nearly all the spacer targets belong to the *Gokushovirinae* sub-family (Figure 4). First identified as phages for intracellular pathogens such as *Chlamydia* and *Spiroplasma, Gokushovirinae* is a very abundant *Microviridae* sub-family found in diverse environments including marine, freshwater, soil, fecal and animal tissue environments (78-90), and in hosts ranging from marine bacteria (65) to enterobacteria (91). Within the *Microviridae*, the only additional hit is to genomes clustering with the proposed *Pichovirinae* sub-family; a single spacer notable mainly because it targets a non-coding region which one would not expect to be well conserved (76) (Fig S2).

**Figure 4.**
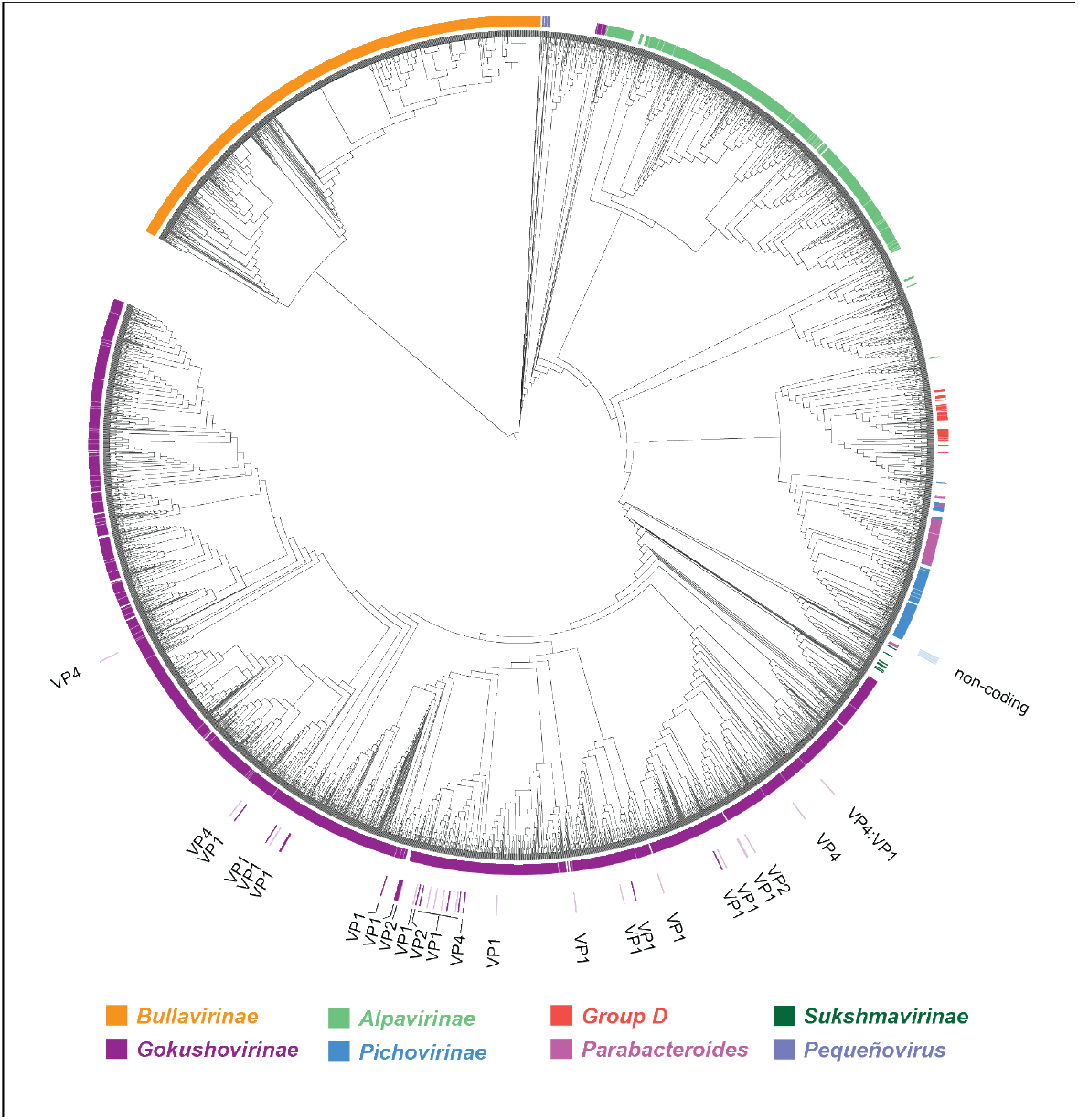
A phylogeny of *Microviridae* capsids reveals the *L. pneumophila* spacer targets belong to the *Gokushovirinae* sub-family. Amino acid sequences of major capsid proteins from 3014 *Microviridae* were aligned using MUSCLE. A phylogeny was generated from this alignment using FastTree and rooted to the *Bullavirinae* clade. Sub-family designations based on the literature are denoted in the inner coloured ring as indicated. Gaps in this ring represent unclassified *Microviridae*. The top hits based on the fewest number of mismatches to *L. pneumophila* spacers are shown in the outer ring, with their target gene designated. VP1 is the major capsid protein, VP2 is the DNA pilot protein and VP4 is the replication initiation protein. Darker spacer bars represent spacers that meet the stringency criteria of 5 or fewer mismatches with its protospacer, the presence of a canonical PAM, and no mismatches in the seed sequence of the spacer.

### Extension of spacer sequence similarity into non-conserved regions of *Gokushovirinae*

We next asked whether one or more of the *Gokushovirinae*-targeting spacers mapped back to regions of low conservation between phages. In particular, spacer sequences that mapped back to predicted host-determinant regions of the capsid (74, 76) might be highly informative, suggestive of a close match. In contrast, highly conserved sequences are present in many related phage genomes, reducing the discriminatory power of an individual spacer to identify the specific phage whose encounter led to its acquisition.

Not surprisingly, the vast majority of *Gokoshovirinae*-targeting spacers map back to regions of high conservation, likely reflective of the overrepresentation of conserved sequences within any database (Figure 5A). Nevertheless, two spacers flank the predicted host-determinant variable region and extend into it (Figure 5B) and one spacer maps to internal variable sequence (Figure 5C). This is potentially quite informative, given the contribution of the capsid variable region to *Gokushovirinae* host range. In short, these spacers likely provide the first glimpse into specific phage sequences required to infect *L. pneumophila*.

**Figure 5.**
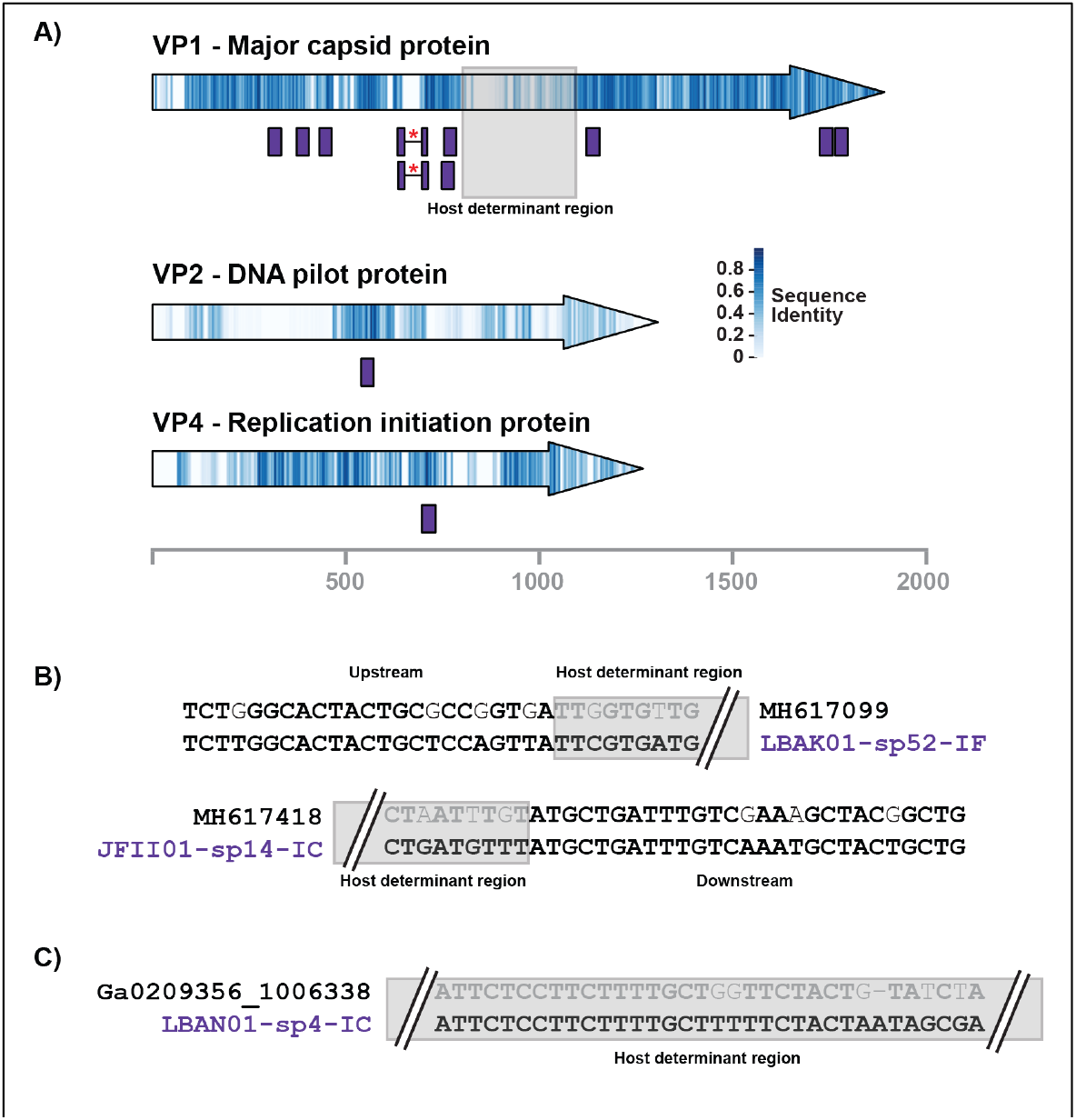
Spacers that meet the stringent criteria largely map to conserved regions of their target genes. **A)** The VP1, VP2 and VP4 gene sequences were aligned for the targeted *Gokushovirinae* genomes that meet the stringency criteria (5 or fewer mismatches with its protospacer, a canonical PAM for its CRISPR subtype, no mismatches in the seed sequence region (positions 1-5, 7, 8)) and are the top hit for a spacer. Nucleotide sequence conservation is shown as a heatmap within the arrow, while spacers mapped onto the alignment are shown as purple boxes. The two boxes marked with a red asterisk contain an insertion in the protospacer region that is present in 1/15 genomes used in the alignment. The host determinant region of the major capsid protein is denoted with a grey box. **B)** Sequence alignments of two spacers that border the host determinant region with their respective top hit phage genome. The spacer name is shown in purple, mismatches between the spacer and the protospacer are non-bolded characters, and the grey box shows the portion of the spacer that is in the host determinant region. **C)** Sequence alignment of a spacer that is contained within the host determinant region. Labelling as in B.

## DISCUSSION

We previously demonstrated that a 30+ kb episomal element, LME-1, is a frequent target of an evolutionarily diverse set of *L. pneumophila* CRISPR-Cas arrays. We followed this observation by demonstrating that these systems actively defend against the transfer of this element between strains (21). We report here that LME-1 represents the only integrated element targeted by *L. pneumophila*’s CRISPR-Cas defenses. Unexpectedly, we also show that *L. pneumophila* CRISPR-Cas defenses are also directed against sequences with strong similarity to the *Gokushovirinae* family of phages. While the specific phages that target *L. pneumophila* remain uncharacterized, *Gokushovirinae* meet the criteria expected for a heretofore undiscovered *L. pneumophila* phage: they are largely lytic, consistent with the dearth of prophage sequences across *Legionellae*; and they are known to infect other intracellular bacteria, such as *Chlamydia* (14-17).

Identification of *L. pneumophila* phages would be of practical importance for at least two reasons. First, once established within water systems, *L. pneumophila* can be notoriously difficult to eliminate. Several studies have observed widespread, decades-long colonization of hospital water systems and stable bacterial genotypes, despite remediation efforts (92-96). A targeted biological remediation strategy (such as phage) would likely pose no threat to human health and could support the frequent treatment of high-risk water systems (to either eliminate or prevent colonization by *Legionellae*) (97). Secondly, the suitability of phage therapy for intracellular pathogens remains an open question. Compared to *Chlamydia, L. pneumophila* is much more genetically tractable, can be cultured outside of host cells in the laboratory, and grows inside a staggering diversity of eukaryotic hosts (4). As such *L. pneumophila* would represent a powerful model system by which to understand the feasibility and constraints of using phages to target intracellular pathogens.

By leveraging the active, adaptive nature of *L. pneumophila* CRISPR-Cas, we have developed an updated, sequence-based composite sketch of *L. pneumophila* phage. Unlike LME-1, which we were able to isolate from defenseless “lysogens,” the *Gokushovirinae* sequences identified in this study are exclusively metagenomic contigs from mixed environmental samples. Our findings suggest that isolation of phages from *Legionella*-containing water sources should be performed under conditions likely to capture gokushoviruses (64, 65, 98), using a diverse collection of CRISPR-mutant strains. The size of gokushoviruses (4-5 kb) also raises the intriguing possibility that synthetic biology could be used to develop one or more *Legionella*-infecting virions (99), starting with the sequences we have identified to date.

During infection, *L. pneumophila* and other intracellular pathogens trade diminished access to nutrients for the protection and isolation provided by the vacuolar environment. Once inside the vacuole, *L. pneumophila* recruits nutrients, avoids lysosomal fusion, and is protected from environmental stresses and host antimicrobial activities (9, 100). Do gokushoviruses access *L. pneumophila* (and *Chlamydia*) inside these “cozy niches?” Is the pathogen’s exposure to phages limited to a transient extracellular period (during cell-to-cell spread) or indicative of a more complicated environmental lifestyle? One thing is clear, 100 years after their discovery, phages continue to hold important secrets about the bacteria upon which they prey.

## ACKNOWLEDGEMENTS

The authors thank Eric Bastien and Melissa Duhaime for early access to their metagenomic data to search for additional *L. pneumophila* CRISPR-Cas targets. In particular, we want to acknowledge the entire community of researchers who have shared data through JGI, especially Karthik Anantharaman, Vincent Denef, Katherine McMahon, David Walsh, and Erica Young for access to their unpublished data. We thank Félix Croteau and Alexander Hynes for their suggestions regarding *Legionella* prophage analysis, Jordan Lin for his help with OrthoMCL analyses and David Faguy for discussions regarding coding, along with members of the Ensminger laboratory for their suggestions and careful reading of the manuscript. SRD is supported by a fellowship from the Department of Biochemistry, University of Toronto and an Ontario Graduate Scholarship. This work was supported by a Project Grant from the Canadian Institutes of Health Research (PHT-148819).

---

**Supplemental Figure 1.**
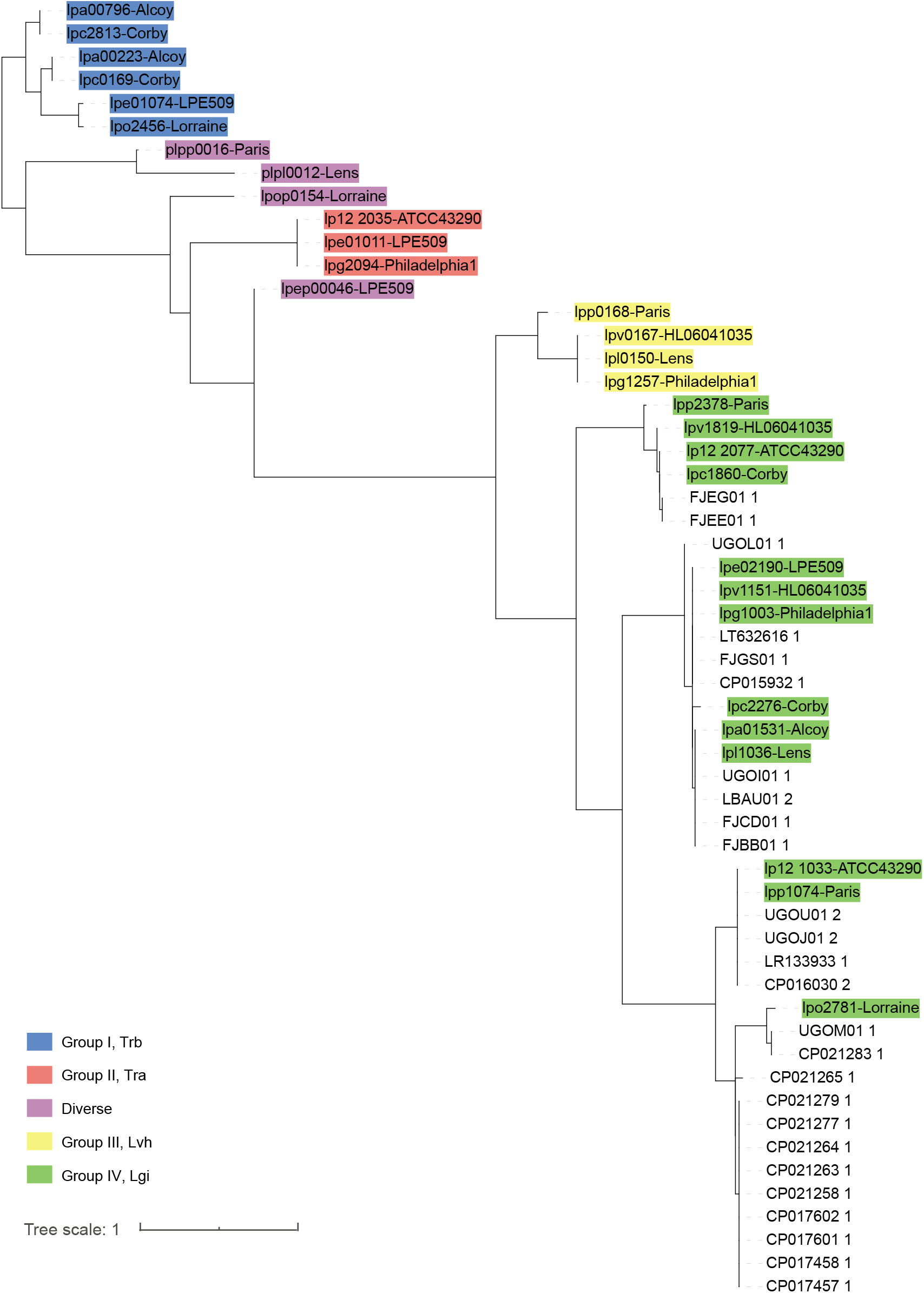
A phylogeny of CsrT sequences reveals mobile elements uncovered by PhiSpy and VirSorter belong to the group IV Lgi elements. CsrT amino acid sequences from mobile elements identified by PhiSpy and VirSorter were extracted and aligned using MUSCLE. A phylogeny was constructed using FastTree and rooted to group I, Trb. Reference sequences were annotated as per Abbott and colleagues (Abbott et al., J Bacteriol. 198, 553–564, 2016) and coloured as indicated.

**Supplemental Figure 2.**
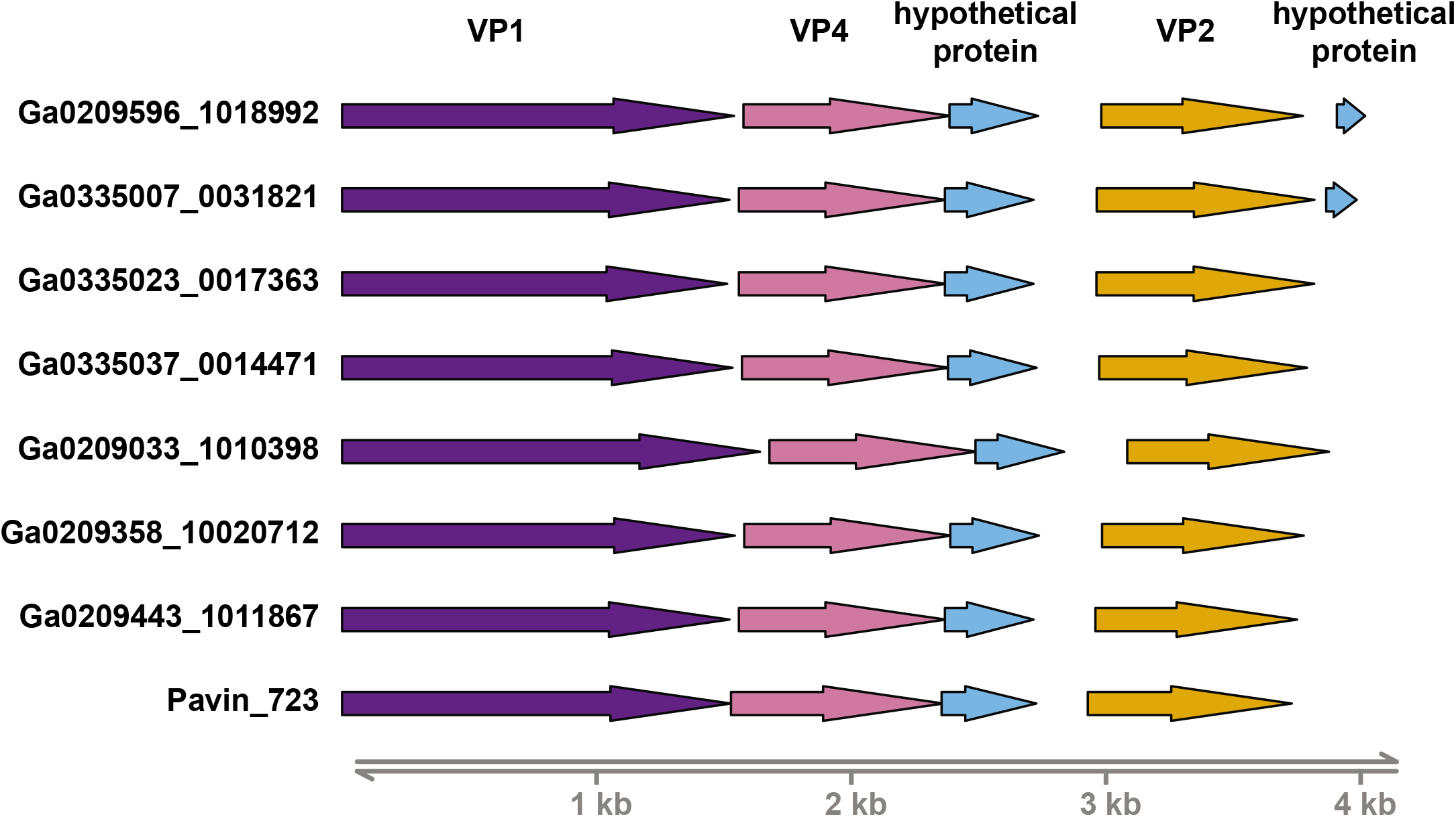
Genome organization of metagenomic contigs that cluster with *Pichovirinae* in the capsid phylogeny. Circularizable metagenomic contigs that cluster with *Pichovirinae* were annotated using Prokka and plotted in GViz with the reference pichovirus Pavin_723. The metagenomic contigs have the unique genome organization VP1-VP4-VP2 characteristic for *Pichovirinae* (S. Roux et al., Plos One. 7, e40418,2012) suggesting that they are indeed part of the *Pichovirinae* sub-family. VP1 (major capsid protein) is shown in purple, VP4 (replication initiation protein) is shown in pink, VP2 (DNA pilot protein) is shown in orange and hypothetical proteins are shown in blue.

**Supplemental Figure 3.**
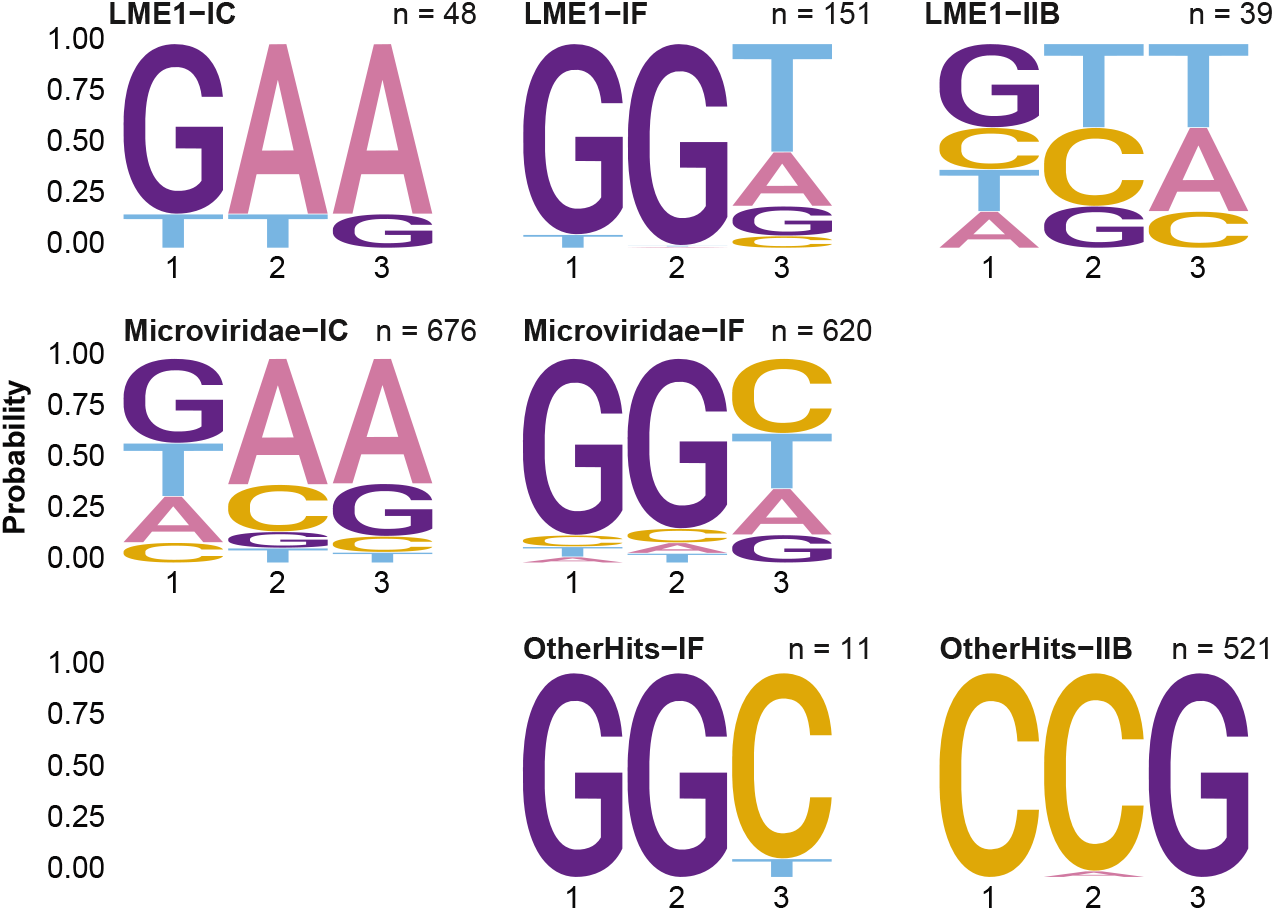
PAM analysis from putative targets reveals that many targets possess the canonical PAM for their CRISPR sub-type. A sequence logo of PAMs from putative targets shows that many targets possess the canonical PAM for their CRISPR sub-type. This suggests spacers were not over-designated to target groups despite being PAM agnostic during data analysis.

